# Assessing multiple evidence streams to decide on confidence for identification of post-translational modifications, within and across data sets

**DOI:** 10.1101/2022.12.15.520504

**Authors:** Oscar M Camacho, Kerry A Ramsbottom, Andrew Collins, Andrew R Jones

## Abstract

Phosphorylation is a post-translational modification of great interest to researchers due to its relevance in many biological processes. LC-MS/MS techniques have enabled high-throughput data acquisition with studies claiming identification and localisation of thousands of phosphosites. The identification and localisation of phosphosites emerge from different analytical pipelines and scoring algorithms, with uncertainty embedded throughout the pipeline. For many pipelines and algorithms, arbitrary thresholding is used, but little is known about the actual global false localisation rate in these studies. Recently, it has been suggested using decoy amino acids to estimate global false localisation rates of phosphosites, amongst the peptide-spectrum matches reported. We here describe a simple pipeline aiming to maximize the information extracted from these studies by objectively collapsing from peptide-spectrum match to peptidoform-site level, as well as combining findings from multiple studies while maintaining track of false localisation rates. We show that the approach is more effective than current processes that use a simpler mechanism for handling phosphosite identification redundancy within and across studies. In our case study using 8 rice phophoproteomics data sets, 6,368 unique sites were identified confidently identified using our decoy approach compared to 4,687 using traditional thresholding in which false localisation rates are unknown.

## Introduction

Phosphorylation is a post-translational modification (PTM) consisting of a phosphate group being bound to one of multiple possible amino acids, including the “canonical” Ser, Thr and Tyr, as well as reports of more rare sites on Arg, His, Cys, Lys and potentially several other residues [1, 2]. The great majority of studies in eukaryotic systems focus on phosphorylation of Ser, Thr, Tyr (STY) amino acids, as the most prevalent and easy to identify [3]. It is now routine to identify (and quantify) large numbers of phosphorylation sites (phosphosites), using LC-MS/MS in data dependent acquisition (DDA) mode, followed by sequence database search. The search algorithm uses the mass/charge measured from precursors i.e. intact peptide sequence (plus PTMs where present) in the MS^1^ scan, and from their fragment ions (in the MS^2^ scan), to compare against theoretical spectra generated computationally from a peptide-sequence database. For analysis of phosphoproteome data (e.g. where samples have been enriched for phosphopeptides), a search would include a parameter for a variable modification, whereby, every STY residue is assessed with and without the addition of the phosphate mass (+79.97 Daltons). There are a large number of MS search engines, including both open and closed-sourced software, for a review of phosphoproteomics pipelines see Locard-Paulet *et al*. 2020 [4] or general phosphoproteomics methods see Riley *et al*. 2016 [5].

Protein modifications may be biological PTMs, or technical artefacts induced during sample handling. Completely unambiguous identification of peptidoforms (a peptide’s amino acid sequence and exact positions of modifications) from LC-MS/MS is often impossible, due to imperfect fragmentation of peptides during MS, multiple peptidoforms in a database sharing some identical or similar sets of fragments ions, near isobaric peptides eluting from the LC column at the same time, causing difficult to interpret chimeric MS^2^ spectra, and multiple other causes. These challenges will lead to some incorrect matches, due to the inferential nature of LC-MS/MS proteomics analyses. There will also be different degrees of confidence among identifications according to how closely observed data matches those extracted from theoretical databases. Hence, analysis pipelines have developed algorithms to calculate scores representing the level of confidence in those identifications being correct [6, 7]. There are two main parts to PTM identification, firstly, a peptide must be “identified” in relation to a reference database, generating a score as a metric of confidence or similarity between precursor ion and candidate peptides in the reference database. Depending on the tool or algorithm these scores might be probabilistic or not [8]. Secondly, once a candidate peptide with n modifications has been identified as the best candidate for a particular spectrum or peptide-spectrum match (PSM), it is common practice in PTM analysis to run another algorithm (within the search engine or as post-processing), which then goes on to identify the n modification within the possible sites m in that peptide, where n<m. In this second step, scores are calculated for all combinations of n modified sites in m possible positions [9, 10].

Some search engines claim these scores to be probabilistic. Truly probabilistic scores would allow unbiased interpretation of scores within and between data sets, for example a score of 0.98 would indicate that PTM to be correct with a confidence of 98%, or among the population of matches yielding a score of 0.98, then 98% would be correct and 2 % incorrect matches independently of the experimental characteristics of the study or data set being analysed. It is difficult to assess whether these scores are truly probabilistic or not, but it has been observed that the same matches obtained from different probabilistic pipelines could have significantly different scores [4] which hinders comparison or combination of PTM data from multiple data sets.

Two well-known PTM localisation scoring algorithms are PTMProphet [7] and ptmRS [10]. PTMProphet uses peak intensities and the number of peaks as parameters in Bayesian mixture models to estimate the probability for each candidate site being modified, before normalising these probabilities according to the number of modified sites in the PSM. While ptmRS utilises the number of characterising ions, those exclusive to the modification, to calculate the probability of each match being random using the hypergeometric distribution. In both cases, PTMProphet and ptmRS, the specificity of the observed data is assessed by comparing the best score for each candidate site versus the next best score for a different site. It is typical practice for PSM false discovery rate (FDR) thresholds to be used to filter most confident peptide identifications (often at 1% FDR) before site localisation scores are generated. For example, users of the Trans-Proteomic Pipeline (TPP) [11] would generally use PeptideProphet [12] to select PSMs based on a pre-defined FDR threshold; this would be followed by PTMProphet scoring which is used to select PTM site localisation or, alternatively, the final score for each site could be a combination of the PSM identification score probability and the localisation score [13]. Simple cut off points of the final PTM localisation scores are used to establish different levels of uncertainty among the PTM identified. The cut offs are usually determined by benchmarking the scores with respect to synthetic data sets for which the key answer is known and hence allow approximate estimation of false localisation rates (FLR). Although there have been attempts to calibrate and demonstrate performance using synthetic data sets [14] it has also been argued whether synthetic data sets are representational of biological variability found in natural data sets [15] and although they are useful means to provide guidance on performance, they are not necessarily an exact indication of how scores could behave on natural data sets. Furthermore, a ptmRS score is a local statistic, giving only information about the confidence in one localization from one PSM. It does not follow that thresholding at a given ptmRS score would deliver particular performance for a global statistic (false localization rate) across the entire data set. Within this context, scores are used as nominal indicators to set arbitrary thresholds following the software developers’ guidance. Given the scores are not likely to be exactly probabilistic and confirmation of identifications via follow-up studies would be too onerous or impossible, it is not known what proportion of sites are being incorrectly identified within the specified threshold. While in peptide identification empirical methods to estimate global statistics are well established, for example based on the inclusion of decoy databases in addition to the target database [16, 17]. In this approach, incorrect matches to the decoy database in a list of PSMs ranked by score, provide FDR estimates that are used to establish more or less stringent thresholds – commonly, trading between the overall number of identifications and the proportion of false positives that the researcher is willing to accept. The proteomics field has largely stabilised at 1% FDR as being acceptable (at the peptide and/or protein level, depending on the type of study).

For PTM analysis, it has been suggested that decoy amino acids should be introduced as well [13, 15]. A decoy amino acid could be any amino acids which cannot naturally be modified by the specified PTM, and thus can be included as a variable modification option. As per peptide identification, matches to decoy amino acids could be useful to provide objective estimates of global FLR in PTM site identification studies. For example, in phosphorylation studies, from several candidates, Ala was chosen as decoy amino acid to generate false positives distributions and estimate FLRs [13]. Ramsbottom *et al*. concluded that including decoy amino acids could enable estimation of global FLR, with the advantageous property that counts of hits to decoy amino acids prove a useful empirical estimate of any ways in which the analysis pipeline might have gone wrong, including both incorrect peptide sequence identification, and identification of incorrect sites within those peptides. However, further research is required to investigate how the primary output from PTM site identification studies using decoy amino acids should be post-processed aiming to optimise the information presented to researchers and facilitate its interpretation, which is the topic of this work. This research assesses core topics in post-processing of PTM sites identification using decoy amino acids leading to the recommendation of a simple post-processing pipeline when using decoys amino acids for PTM site identification.

The first topic relates to the cumulative evidence for sites within a data set. The primary output from statistical processing is local scores or statistics for a PSM, which can then be collated to a data set level (global), using matches to decoy peptides and decoy amino acids to assess global FDR/FLR on the set of PSMs supporting PTM sites. For abundant peptidoforms, which here we define as the peptide sequence plus the exact set of modifications carried on exact residue positions, it is common for them to be sampled multiple times by the instrument in different scans, leading to redundant PSMs reporting on the same peptidoform. A research team would generally wish to “collapse” the redundancy and re-calculate statistics, so that each peptideoform is reported only once, even if it was supported by more than one PSM. Common practice is simply to take the highest scoring PSM or highest localisation score (following say thresholding PSMs first at 1% FDR). However, this has the clear disadvantage of information loss; it seems intuitive that peptidoforms supported by multiple PSMs are more likely to be true by those supported by fewer. We wish to develop methods to explore, understand and handle this phenomenon appropriately, such that phosphosite and PSM count can be taken into account.

The next step we wish to address concerns appropriate handling of how PTM site information should be collapsed from PSM to peptidoform level. Traditionally, for each peptidoform, researchers have simply selected the maximum score within all PSM-site candidates matching to the peptidoform. However, it has not been assessed whether scores for different PTM sites within a PSM should be considered independent from each other (then taking the maximum would be appropriate) or they are not and, a different statistic across the PSM would be preferable.

Finally, there is much to be gained when identifying PTMs on a large scale by combining results from multiple studies in a meta-analysis. Particularly, we assess compatibility of scores from independent analyses and how information about a site being identified in several independent analyses could be used to increase our confidence on those sites’ localisation being correct.

## Methods

### Data sets

The post-processing analyses are illustrated using 12 different data sets. Two are “synthetic” data sets, which were manufactured to contain specific peptidoforms, and thus can be used to assess algorithm performance with a known answer. The other 10 belong to three species: 1 for Arabidopsis thaliana, 1 Human (both data sets explored in [13]) and 8 rice data sets (data sets used to create a comprehensive meta-analysis of the rice phosphoproteome, manuscript in preparation). It is expected that the variety of data sets would allow methodology performance assessment while using the synthetic data sets and investigate whether similar patterns would be observed in biological data sets, between and within species. The full list of data sets with a brief description of the experimental objectives extracted from their publications’ abstracts can be found in Table 1. Reference databases used for PTM searches are also listed in Table 1.

**Table 1.** Data sets used to illustrate analyses with a brief description of the experiments performed and files selected for analyses.

| Data set<br>ProteomeXchange<br>Unique ID | Species | Reference<br>Database | Brief description | References |
| --- | --- | --- | --- | --- |
| PXD000138 | Synthetic | The source database containing IPI human sequences and phosphopeptide libraries (matched to the original study) | 96 synthetic tryptic peptide libraries, analysed using LC-MS/MS on an Orbitrap mass spectrometer with either HCD or ETD as fragmentation technique. 10 files were used for this analysis. | [18] |
| PXD007058 | Synthetic | Library formed by synthetic peptides | Files named “HCDOT” (i.e. HCD mode on an Orbitrap) pools 1 to 5, reps 1 and 2 were used. | [14] |
| PXD008355 | *Arabidopsis thaliana* | Araport11 sequences | Proteome-wide view of the plant target of rapamycin phosphorylation and interaction landscape using 4 biological repeats. 12 files were used for this analysis. | [19] |
| PXD000612 | Human | Created from the Level 1 PeptideAtlas Tiered Human Integrated Search Proteome, containing core isoforms from neXtProt | HeLa S3 cells untreated, mitotically arrested and released, or stimulated with epidermal growth factor (EGF) and additionally, they performed immunoaffinity enrichment of Tyr phosphorylated peptides from untreated, mitotic, EGF-stimulated, and pervanadate-treated (for PTP inhibition) cells. 6 randomly selected files were used. | [20] |
| PXD000923 | Rice: *Oryza sativa* | Built from protein sequences derived from the MSU Rice Genome Annotation Project, the Rice Annotation Project | Study on mature pistil of rice. Using IMAC enrichment, hydrophilic interaction chromatography fraction and high-accuracy MS instrument (TripleTOF 5600). | [21] |
|  |  | Database (RAP-DB), including both translated CDS and predicted sequences, and Uniprot, including both reviewed and unreviewed sequences. * |  |  |
| PXD002222 | Rice: *Oryza sativa* | Same as * | Large scale enrichment of phosphopeptides and identification of phosphosites in rice before and 24 h after Xoo infection. | [22] |
| PXD002756 | Rice: *Oryza sativa* | Same as * | Phosphoproteomic analyses for developing rice (*Oryza sativa*) anthers around the time of meiosis. | [23] |
| PXD004705 | Rice: *Oryza sativa* | Same as * | Profiling of rice young seedling phosphosites, phosphopeptides and phosphorylation intensity dynamics in response to ABA. | [24] |
| PXD004939 | Rice: *Oryza sativa* | Same as * | Profiling of rice young seedling phosphosites, phosphopeptides and phosphorylation intensity dynamics in response to ABA | [25] |
| PXD005241 | Rice: *Oryza sativa* | Same as * | Profiling of 6 tissues, including callus, leaves, roots, shoot meristem, young panicles and mature panicles. | [26] |
| PXD012764 | Rice: *Oryza sativa* | Same as * | Phosphorylation is as response to Pi starvation in rice root. | [27] |
| PXD019291 | Rice: *Oryza sativa* | Same as * | Analysis of a rice male-sterile mutant ap1, which produces non-viable pollen grains with defective starch accumulation. | [28] |

### PTM identification and localisation

An analysis pipeline was set up for analysis as described by [13], using the Trans-Proteomic Pipeline (TPP) [11], including the Comet search engine [29], ThermoRawFileParser [30], and post-processing via PeptideProphet [12], iProphet [31] and PTMProphet [7]. The analysis parameters for TPP are displayed in Supp Table 1. The only difference between Decoy searches (pASTY) and a typical search (pSTY) was the inclusion of Alanine as potential phosphorylation site for pASTY searches, the rest of parameters remained the same.

Proteome Discoverer (PD, Thermo Fisher Scientific) was also used, including the Mascot search engine [32], Percolator [33] and the ptmRS site localisation [10, 32]. The PSM probability values were calculated using 1-PEP values (reported natively by the pipeline), with the PTM probability being calculated innately though ptmRS scoring. The analysis parameters for Mascot and ptmRS are shown in Supp Table 2.

Peptide identifications from PeptideProphet scoring were initially filtered at 1% FDR (PSM level), based on the target-decoy search results provided by this tool. Probabilities of potential modified sites in identified peptides were computed using PTMProphet. The final score for each site before post-processing was calculated as (Peptide identification score)*(Site identification score).

### Definitions

pSTY: typical phosphorylation site search looking for a phosphate group at S, T and Y.

pASTY: phosphorylation search including S, T and Y as target amino acids and Alanine as decoy amino acid.

Peptide spectrum match (PSM): peptide sequence with or without modifications matching to a single spectrum.

pAla: is a match to the decoy amino acid Alanine. pAla hits can be present at any reporting level.

PSM-site reporting: at this level of reporting, the results include the scores relating to each PTM site for each PSM passing the peptide identification threshold of 1% FLR. For each PSM there will be as many PSM-sites as phosphosites identified in the peptide. At this level there is duplication of identical matches to sequences and sites, as there could be multiple PSMs supporting the same peptidoform.

Peptidoform: this is a unique peptide in the data set formed from the combination of peptide sequence and the number and position of modifications identified.

Peptidoform-site reporting: this concept refers to results reporting each specific phosphosite within each peptidoform. At this level, duplication of identically-modified sequences have been removed but all unique sequences and sites to target and decoy matches identified at PSM-site level still remain in the data set.

### Post-processing

Data analyses were performed in R 4.0.3 via RStudio Version 1.1.442 (© 2009-2018 RStudio, Inc.). The following steps were used to assess different approaches for post-processing phosphorylation site scores:

#### Step 1: Using phosphosite frequency to inform scores

Under the hypothesis that modified sites that have been observed more frequently in one data set are more likely to be correct than those which have been observed less often, or in other words, random PTM matches are expected to be observed less frequently than matches to correct sites, scores can be modified to include information about how often sites are observed in a data set.

We could consider each phosphorylated site as dichotomous events in which the site can be phosphorylated or not. Based on matches to decoy amino acids the probability of obtaining a random match can be estimated and scores adjusted based on how many times a site has been observed phosphorylated over the possible chances of being phosphorylated. Using this information, scores for all sites can be penalised, adjusting our confidence in those identifications. This approach takes into account differences in PSM counts. We implemented this approach using the binomial distribution which follows the discrete distribution:

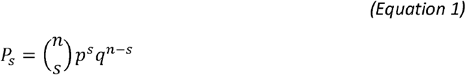

Where *p* is the probability of obtaining in a spectrum a match to pAla by chance (unique pAla count /unique spectrum), *q* is equal to 1-*p*, *s* is the number of successes or frequency in which a specific protein site has been observed phosphorylated across the data set, and *n* is the number of times that same protein position is seen across all PSMs in a data set (phosphorylated or not). This binomial probability can be interpreted as a measure of confidence that the site is a true phosphosite given that it has been observed *s* times phosphorylated over *n* possible opportunities.

The final adjusted probabilistic score incorporates this information by simply multiplying the site score by its binomial score (1-*P_s_*) for each PSM-site. This approach will be referred to as Binomial_Adjustment.

#### Step 2: Collapsing from PSM-site to Peptidoform-site level

Collapsing from PSM-site level to peptidoform-site level is most commonly achieved by simply taking the maximum final site score (or probability) within all PSMs with identical sequence and position being phosphorylated. This approach considers the maximum to be the best representation of its class and assumes that scores belonging to the same PSM are independent from each other. Chosen scores for a peptidoform may come from different PSMs. This form of collapse was produced for unadjusted data and data after Binomial_Adjustment, results will be referred as Unadjusted_PformMax and Binomial_PformMax respectively.

Another potential approach we tested for collapsing results from PSM-site level to peptidoform-site level was taking the mean score of the PSM-sites belonging to a peptidoform-site. In this approach the true probabilistic score would be the centre of the distribution formed by all PSM-site scores linked to the peptidoform-site. As for the maximum approach, PSM-site scores would be related to other PSM-site scores for the same peptidoform-site but independent from any other site. These results will be referred as Unadjusted_PformMean and Binomial_PformMean for unadjusted and binomial adjusted PSM data.

In a third approach, we tested using the mean score of the phosphosites in a PSM and then taking the maximum mean score for all PSMs linked to the same peptidoform i.e. the PSM delivering the best overall explanation for the sites scored. This approach considers the scores from a PSM to be dependent from each other which would make a PSM the basic experimental unit and not the PSM-site. Results from this collapse method will be referred as Unadjusted_PformMM and Binomial_PformMM.

Finally, considering PSMs for the same site independent from each other (and independent from other sites within a PSM), the product of (1 – (site probability)) could be understood as cumulative information of a modified site being correctly identified given the observed data. This approach would assume that each observation of a site is independent of others (an assumption we know is unlikely to be true). This will be referenced later in the report as Product_Pform.

An example of results collapsing from PSM-site level to peptidoform level are displayed in Figure 1.

**Figure 1.**
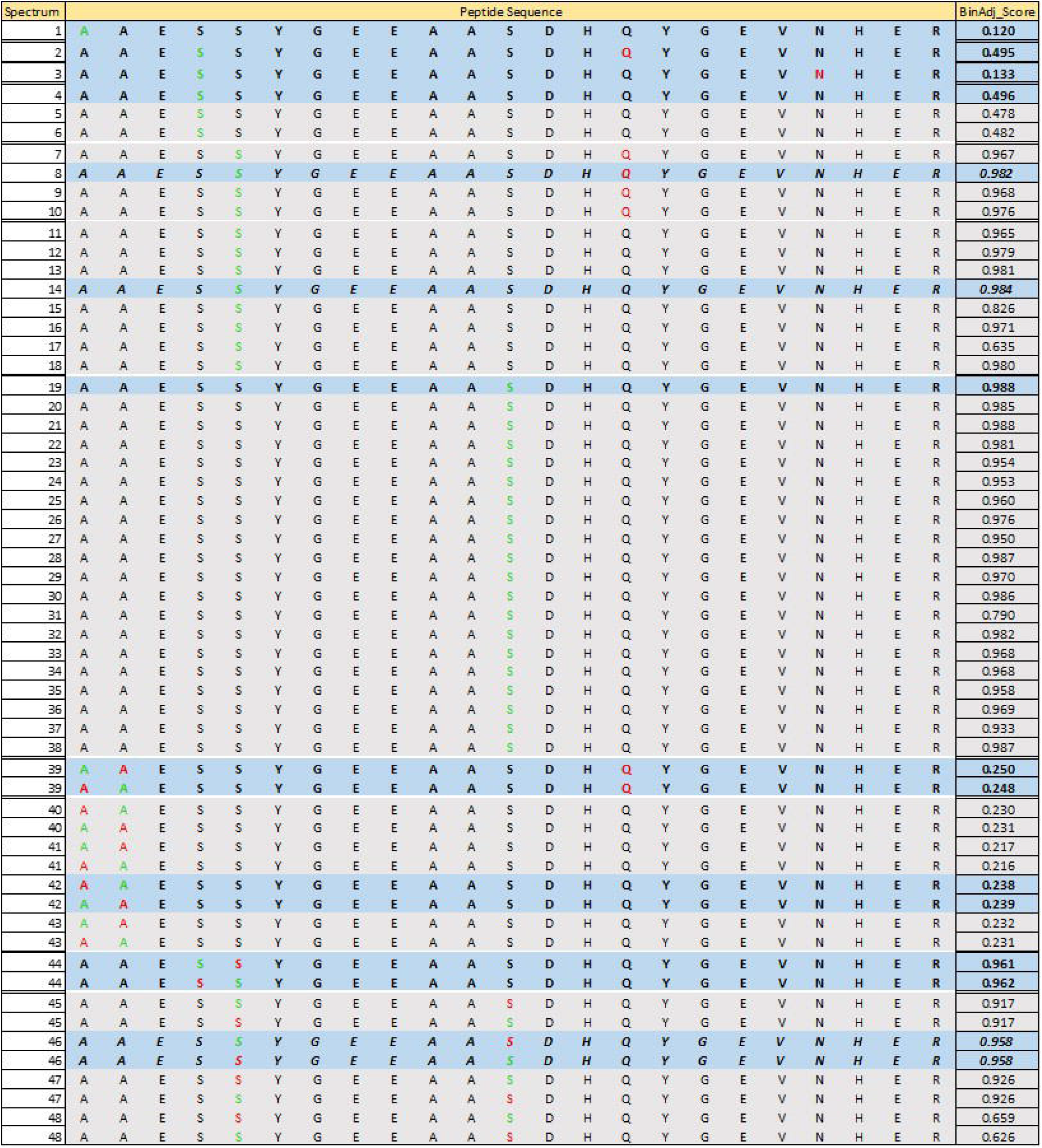
Illustration of results at PSM-site level for a peptide-sequence from the Arabidopsis data set. Phosphosites being scored are coloured in green while any other modification sites identified in the spectrum are coloured in red. BinAdj_Score indicate that these are scores after binomial adjustment. The Peptidoform-site level observations which will result after taking the maximum are in bold and highlighted in blue.

For adjusted and unadjusted data FLR, based on decoy matches, the FLR formula from Ramsbottom *et al*. [13] was adapted as:

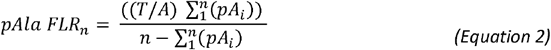

*T* is the total count of target amino acids (STY), *A* is the total count of decoy amino acid in the data set, 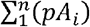 is the count of phosphorylated decoy amino acid to the *n* observation in the data set and *n* is the count of observations at a given position in the decreasing ranked list of scores. We took a conservative approach when calculating the pAla FLR by considering decoys, all sites in a PSM with a decoy match. We have also adapted the FLR calculation from (13). In that publication, FLR is defined as the false localisations amongst both the known wrong hits i.e. pAla sites, and the within the STYs, with the following formula

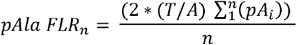

The formula we use in Equation 2, assumes that pAla results are ultimately removed (hence the subtraction term in the demoninator), and we do not multiply the numerator by a factor of 2 to account for (silent) false positives with STYs and the known false positive pAla sites. We believe that in practice the formula reported in (13) gives highly conservative results, and for the vast majority of uses of the data, it is more natural to remove pAla hits at the final reporting level, as known false positives.

Note that for all FLR calculations, there might be ties between scores for multiple sites and therefore the order of those sites may influence the results, specifically if there are decoy sites among the equally scoring sites. Any kind of ordering based on the amino acids being phosphorylated could introduce bias by placing decoy amino acids at the beginning or end of equally scoring sites and hence, it could over or underestimate FLR. To promote randomness, we just ordered our results alphabetically in relation to the amino acid sequence but any other ordering not related to the amino acid being modified should yield similar results.

For synthetic data sets, FLR were calculated based on the key answer from these studies as:

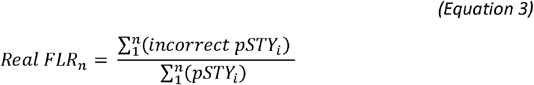

## Results

### Using phosphosites frequency to inform scores

Given the premise that random incorrect PTMs happen less frequently than correct matches, information about the number of matches to each site observed in the data set could be used to modify our confidence of each site being correct. Figure 2 shows this to be the case for all 10 natural data sets used in this report. Although there are some decoy matches (pAla) which can be observed up to 100 times i.e. PSM count, the counts of PSMs supporting target amino acids are consistently higher than that for decoys (10 top panels). Zooming to those cases with fewer than 50 observations allow us to clearly see higher counts of PSM matches and higher medians for target matches than decoy matches (10 middle panels). This effect is emphasised if only those sites with probability above 0.95 are included in the graph (10 bottom panels). Similar patterns can be observed for results from the PD pipeline (Supp Figure 1). We can thus conclude that PSM count supporting a PTM site identification is informative, and that taking a maximum PTM score to collapse redundancy is likely to lose valuable information, as a PTM site, for example, with score = 0.97 with 1 observation, or with 100 observations would score equally after collapse, but are not equally likely to be true.

**Figure 2.**
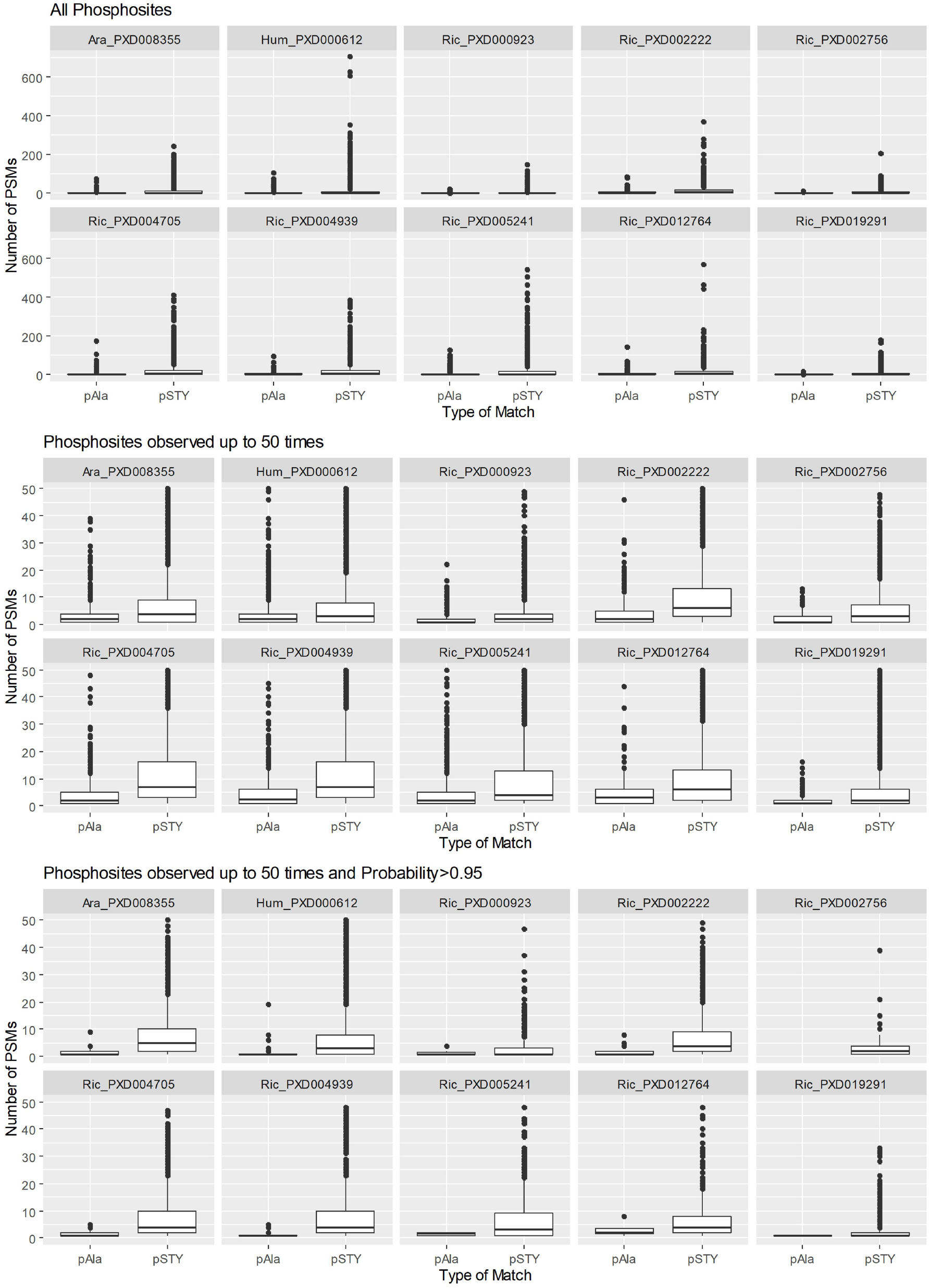
Number of PSMs matching specific sites by type of match for human and Arabidopsis natural data sets analysed using the TPP pipeline. pAla matches correspond to decoy phosphosites and pSTY target matches. Top 10 panels display all data, middle 10 panels display phosphosites observed up to 50 times and, the bottom 10 panels are those phosphosites with scores probability above 0.95 and observed up to 50 times.

In the Binomial_Adjustment approach, each phosphosite is considered to be a dichotomous outcome where the site can be phosphorylated or not. Based on the binomial distribution, we calculated the probability of each phosphosite being random given that it has been observed phosphorylated s times over the n times that site occured in the data set, independently of being phosphorylated or not. Then, the complement of these probabilities can be used to penalise the unadjusted scores for all sites, penalizing more heavily those that have been observed less often. We used two natural data sets (Ara_PXD008355 and Hum_PXD000612) and two synthetic (Syn_PXD000138 and Syn_PXD007058) to make those comparisons based on the pAla FLR (Figure 3). In Figure 3, especially in the Arabidopsis data set, it can be clearly observed the effect of high scoring random matches with two small peaks at low FLR values for the unadjusted scores (blue). A successful adjustment method would smooth those early peaks by pushing incorrect random matches down the list. The Binomial_Adjustment method has the desired effect across all data sets, with a smoothing effect resulting in local FLR estimates staying below or at a comparable level to the FLR from unadjusted results (Figure 3). This adjustment takes into account differences in peptide abundance by considering how many times a particular site has been observed phosphorylated or not.

**Figure 3.**
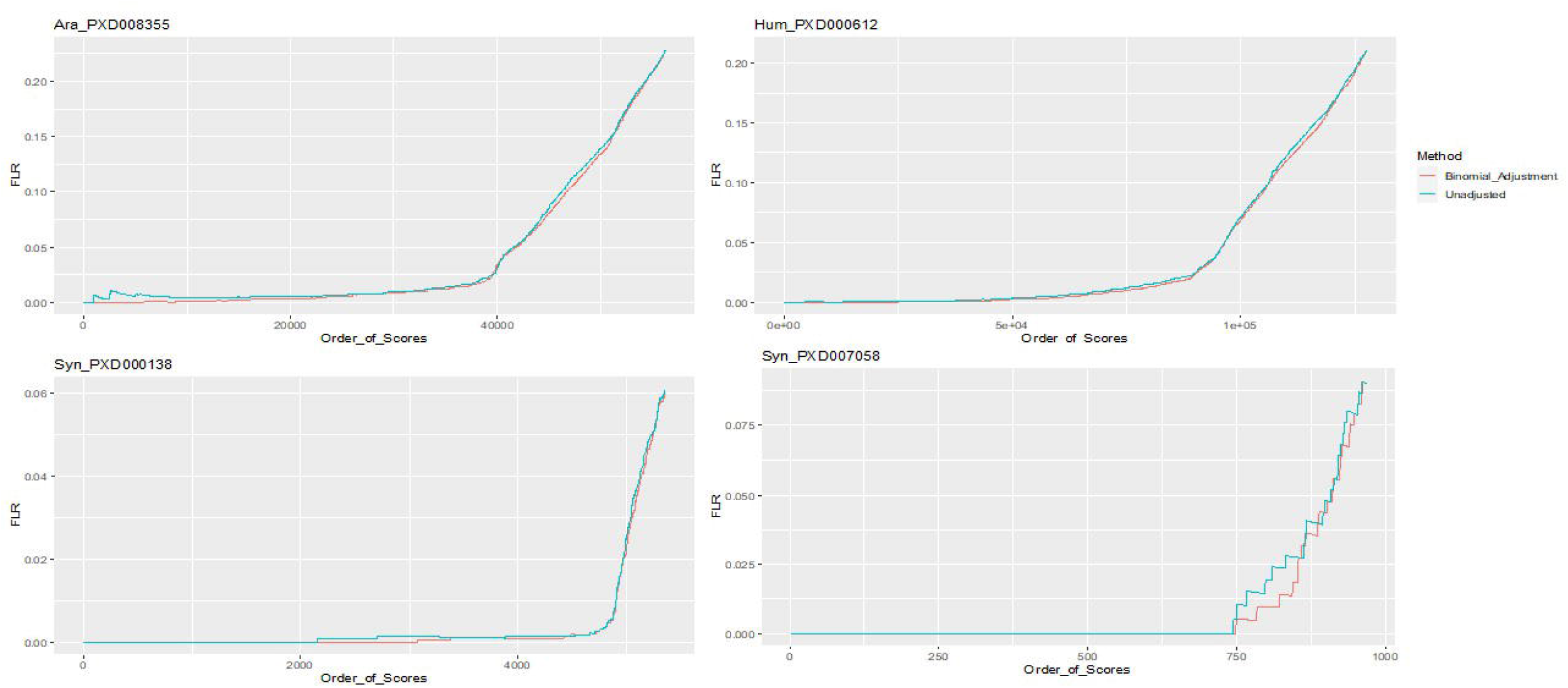
pAla FLR at PSM-site level by ordered unadjusted scores (Blue), using the Product_Adjustment (Green) and, using the Binomial_Adjustment (Red). Data sets analysed using TPP.

The Binomial_Adjustment is expected to produce more accurate estimates at low FLR levels than unadjusted results, before the adjusted and unadjusted FLR convergence is achieved. For these four data sets, at PSM Level, local FLR reached early convergence between the Binomial_Adjustment and unadjusted approaches for all four data sets (Table 2). Table 2 also shows comparability between the Binomial_Adjusted results and unadjusted data in terms of Real FLR, calculated as the proportion of incorrect matches based on the answer key for the synthetic data sets for which the true phosphosites are known.

**Table 2.** Number of PSM-site matches for Arabidopsis, Human and two synthetic data sets analysed using the TPP pipeline and thresholds at 1%, 2.5%, 5% and 10% pAla FLR based on unadjusted results and adjusted using the Product_Adjusted and Binomial Adjusted approaches. For the synthetic data sets, the Real_FLR is also displayed at each pAla FLR threshold.

|  | Arabidopsis PXD008355 |  |  |  |  | Synthetic PXD000138 |  |  |  |  |  |  |  |  |
| --- | --- | --- | --- | --- | --- | --- | --- | --- | --- | --- | --- | --- | --- | --- |
|  | pAla FLR |  |  |  |  | pAla FLR |  |  |  |  |  |  |  |  |
|  | Total | 10% | 5% | 2.5% | 1% | Total | 10% | 5% | 2.5% | 1% | 10% | 5% | 2.5% | 1% |
|  | Number of observations |  |  |  |  | Number of observations |  |  |  |  | Real FLR |  |  |  |
| Unadjusted | 56276 | 46191 | 41619 | 39570 | 31249 | 5360 | 5360 | 5220 | 4997 | 4911 | 5.1% | 3.6% | 2.0% | 1.3% |
| Binomial_Adjusted |  | 46660 | 41799 | 39564 | 32620 |  | 5360 | 5247 | 5013 | 4900 | 5.1% | 3.7% | 1.7% | 1.2% |
|  | Human PXD000612 |  |  |  |  | Synthetic PXD007058 |  |  |  |  |  |  |  |  |
|  | pAla FLR |  |  |  |  | pAla FLR |  |  |  |  |  |  |  |  |
|  | Total | 10% | 5% | 2.5% | 1% | Total | 10% | 5% | 2.5% | 1% | 10% | 5% | 2.5% | 1% |
|  | Number of observations |  |  |  |  | Number of observations |  |  |  |  | Real FLR |  |  |  |
| Unadjusted | 127565 | 106028 | 96406 | 90032 | 70893 | 969 | 969 | 908 | 829 | 748 | 10.9% | 7.5% | 4.1% | 1.3% |
| Binomial_Adjusted |  | 106165 | 96400 | 90289 | 74949 |  | 969 | 911 | 854 | 822 | 10.9% | 7.5% | 4.5% | 3.6% |

The PD pipeline seems to be more successful than TPP avoiding early peaks (Supp Figure 2). The results after using the binomial adjustment appear to also improve at 1% FLR with respect to unadjusted (Supp Table 2).

### Collapsing from PSM-site to Peptidoform-site level

Results at PSM level present many redundancies which could hinder their interpretation. Variations in scoring among equally modified peptide candidates are due to differences in ion fragmentation indicating a higher or lower level of confidence of the site identified. To facilitate interpretation, results can be collapsed at peptidoform-site level as they represent the most basic unique unit emerging from the combination of peptide sequences, the type and number of modifications, as well as the position of those modifications. We investigated three different ways for collapsing the results from PSM to peptidoform level and we applied them to unadjusted results and results after Binomial_Adjustment. Additionally, we collapsed to Product_Pform results by simply multiplying (1-prob) across PSM-sites corresponding to a same protein-site and then, removing redundancies in the data sets. This leads to the seven approaches explained in the methods section and with the pAla FLR curve generated using these methods displayed in Figure 4. The same four previously analysed data sets (two natural and two synthetic) were also used here to investigate the effect of collapsing at peptidoform level.

**Figure 4.**
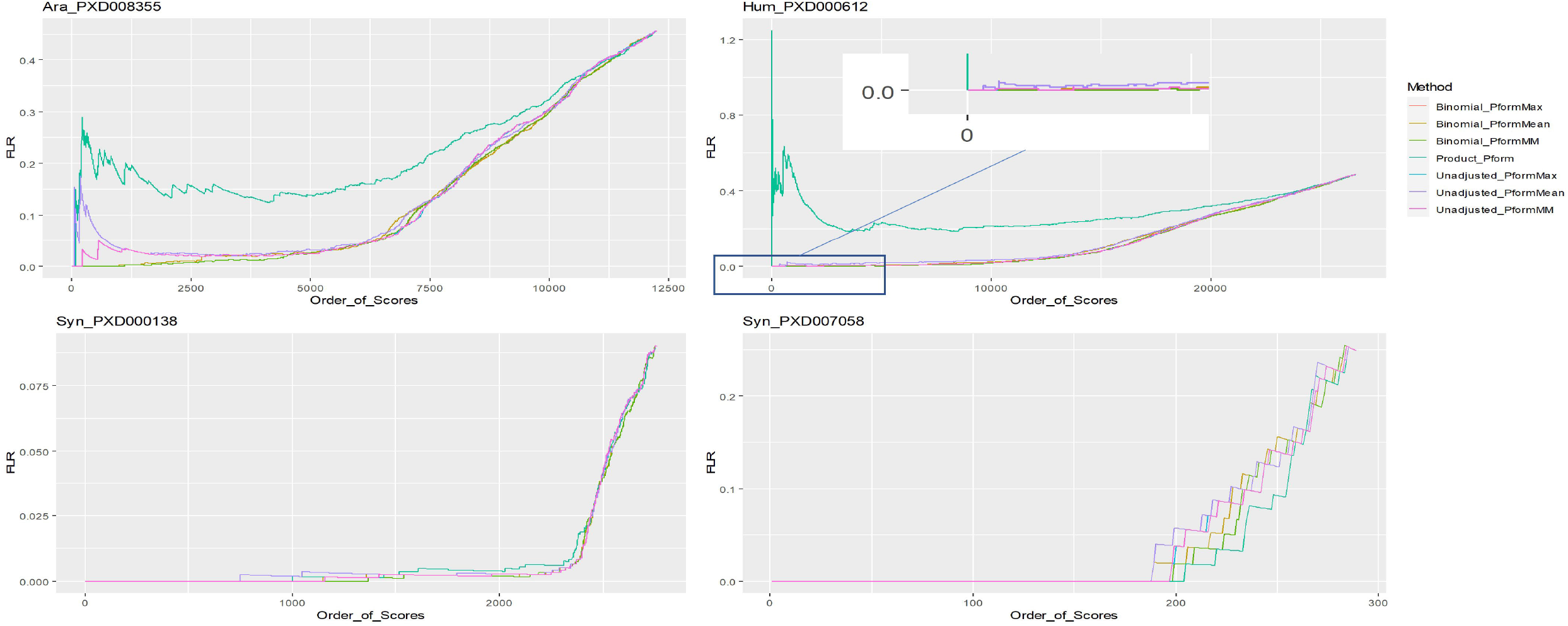
pAla FLR at Peptidoform-site level by ordered scores for the seven collapsing approaches in two natural data sets and two synthetic. Data analysed using TPP.

Clearly, the Product Adjustment does not achieve the desired results (Figure 4). For the natural data sets, the pAla FLR from the Product Adjustment remains much higher than unadjusted results (for an equivalent position in the ranked list) and therefore was confirmed not to be a suitable adjustment and was not included in further analyses. In the same natural data sets, in Figure 4, it can be observed a general better performance of data after Binomial_Adjustment, compared to unadjusted data which exhibit significant early peaks in the pAla FLR curve, indicating high scoring incorrect matches. Binomial_Adjustment data appear to perform better than unadjusted, independently of the binomial collapse method used. Results indicate that collapsing at peptidoform level could emphasise the effect of random incorrect high scoring matches (observed in unadjusted data in Figure 4), but this effect could be successfully mitigated by applying the Binomial_Adjustment.

Amongst the collapsing methods applied after Binomial_Adjustment, the Pform_Mean method appears to underperform compared to the Pform_Max and Pform_MM approaches. It could be expected that not all the distributions of scores for all sites would be centred or well characterised given that for most sites there are only few observations per site in the data sets. Thus, we concluded that the site score mean is not a suitable statistic to represent peptidoform-site scores.

A very similar performance can be observed between Pform_Max and Pform_MM approaches for both binomial adjusted and unadjusted data across all data sets (Table 3). The Pform_Max method assumes that the maximum score for a site is the best representation of that site in the data set. While in the Pform_MM method, all site scores within a PSM depend on the other site scores. Or in other words, the PSM, and not its sites, is considered the basic independent unit. However, if these two methods yield highly correlated results, it would mean that the maximum score in a PSM-site is likely to yield high scores in any other phospho sites within the same PSM. To assess this hypothesis, we calculated the Spearman correlation coefficients, for the four data sets assessed in Table 3, based on the ranked scores between Pform_Max and Pform_MM in ranges of (0.9994 to 1) and (0.9992616 to 1) for binomial adjusted and unadjusted data respectively. While between Pform_Max and Pform_Mean were lower, in ranges of (0.8883875 to 0.9574074) and (0.8536769 to 0.9521967) for binomial adjusted and unadjusted data. For the PD pipeline, correlation between Pform_Max and Pform_MM in ranges of (0.9365 to 1) and (0.9357 to 0.9949) for binomial adjusted and unadjusted data respectively, and (0.8316 to 0.9159) and (0.8439 to 0.9155) for binomial adjusted and unadjusted data. In conclusion, results in Table 3 suggest the Binomial_PformMax and Binomial_PformMM to be the best performing methods and, are highly correlated and thus similar to each other.

**Table 3.**
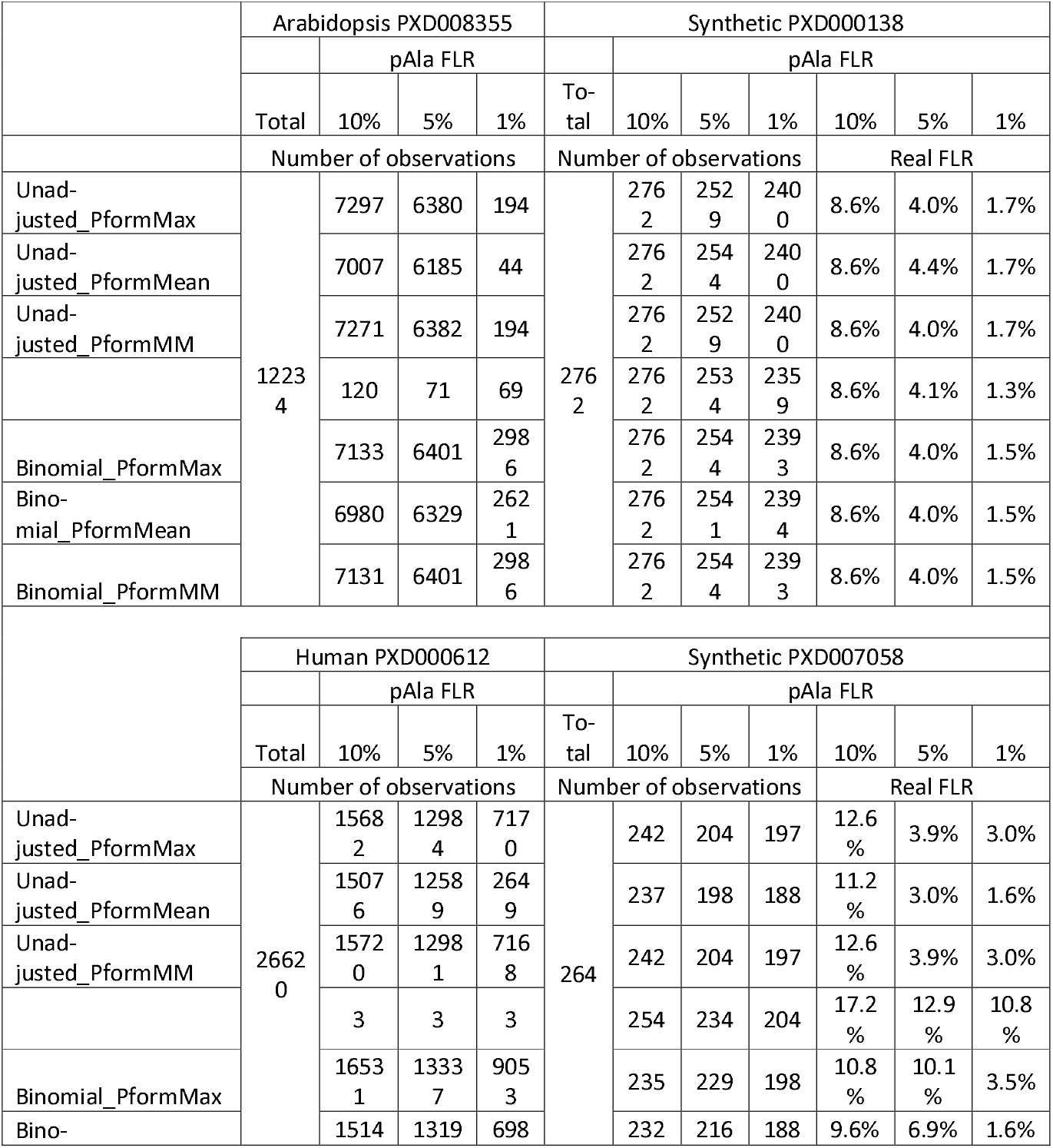
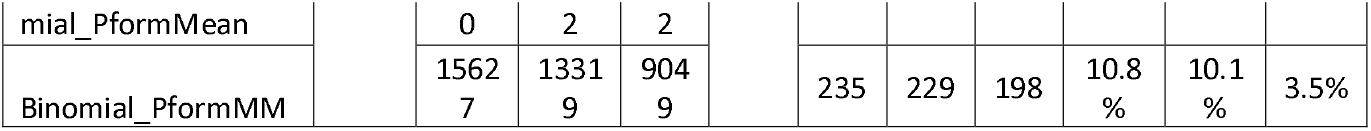
Data sets analysed using the TPP pipeline. Number of phosphosites at Peptidoform-site level for seven collapsing approaches. Results are reported overall and for pAla 1%, 5% and 10% FLR thresholds. Real FLR is also displayed for the synthetic data sets.

For the PD pipeline, the PformMM clearly outperforms any of the other approaches, especially in the natural data sets (Supp Figure 3 and Supp Table 4). Binomial Adjustment appears to have little effect on PD output, similar patterns can be observed between the unadjusted and adjusted data (Supp Figure 3). Comparing results at peptidoform level, the PD pipeline returns a larger number of sites than the TPP pipeline for the natural data sets at all pAla FLR levels. For the largest synthetic data set both pipelines return similar results while for the small data set the performance of PD is very poor.

Considering the limited number of data sets in Table 3, further analyses were performed only using PformMax and PformMM collapsing methods in the 8 rice data sets, as shown in Table 4. Overall, when assessing the total number of sites being identified across all 8 data sets, the binomial adjusted methods appear to perform better than unadjusted methods with more sites being observed after binomial adjustment at either pAla FLR threshold. At the same time, the Binomial_PformMM methods yields more phospho sites than just taking the maximum score (Binomial_PformMax). The improvement in performance seems to confirm the PSM as minimum basic independent unit in the data set and not the PSM-site. Therefore, for post-processing results where PSM-site scores within a PSM are not independent, the mean of the scores within a PSM would be a more appropriate statistic than the individual PSM-sites scores. Drawing our attention to specific data sets performance, the results suggest that PXD002222 and perhaps PXD002756 do not seem to perform as well as the others after binomial adjustment. The observed effect could be due to lack of specificity in the analysis. As shown in Figure 5, scores for both target and decoy matches in PXD002756 seem to follow overlapping distributions, suggesting high level of randomness in PTM localisation and hence have a similar frequency of correct and incorrect matches. To a lower extent, this same effect can also be observed for PXD005241. While for PDX002222, a great proportion of decoy matches achieve high scores, reducing overall discrimination between target and decoy matches.

**Table 4.** Number of phosphosites at Peptidoform-site level for 8 rice data sets. Results are reported overall and for pAla 1% and 5% FLR thresholds. Results are displayed for PformMax and PformMM collapsing approaches from unadjusted and binomial adjusted data.

| Data set | PSM | Peptidoform | Unadjusted_PformMax |  | Unadjusted_PformMM |  | Binomial_PformMax |  | Binomial_PformMM |  |
| --- | --- | --- | --- | --- | --- | --- | --- | --- | --- | --- |
|  |  |  | pAla FLR < 5% | pAla FLR < 1% | pAla FLR < 5% | pAla FLR < 1% | pAla FLR < 5% | pAla FLR < 1% | pAla FLR < 5% | pAla FLR < 1% |
| PXD000923 | 10770 | 5340 | 3120 | 716 | 3119 | 716 | 3165 | 1588 | 3169 | 1588 |
| PXD002222 | 26063 | 5380 | 2430 | 1293 | 2429 | 1293 | 2521 | 1200 | 2540 | 1200 |
| PXD002756 | 13565 | 6972 | 642 | 300 | 644 | 300 | 724 | 281 | 725 | 281 |
| PXD004705 | 56946 | 8782 | 3428 | 489 | 3425 | 489 | 3433 | 1760 | 3450 | 1727 |
| PXD004939 | 56078 | 8650 | 3463 | 1796 | 3475 | 1796 | 3496 | 1720 | 3498 | 1710 |
| PXD005241 | 81152 | 19107 | 792 | 186 | 798 | 186 | 781 | 324 | 788 | 324 |
| PXD012764 | 18535 | 5241 | 1275 | 87 | 1272 | 87 | 1246 | 897 | 1244 | 897 |
| PXD019291 | 28472 | 18054 | 10058 | 4646 | 10392 | 4662 | 11126 | 5510 | 11375 | 5509 |
| ALL | 291581 | 77526 | 25208 | 9513 | 25554 | 9529 | 26492 | 13280 | 26789 | 13236 |

**Figure 5.**
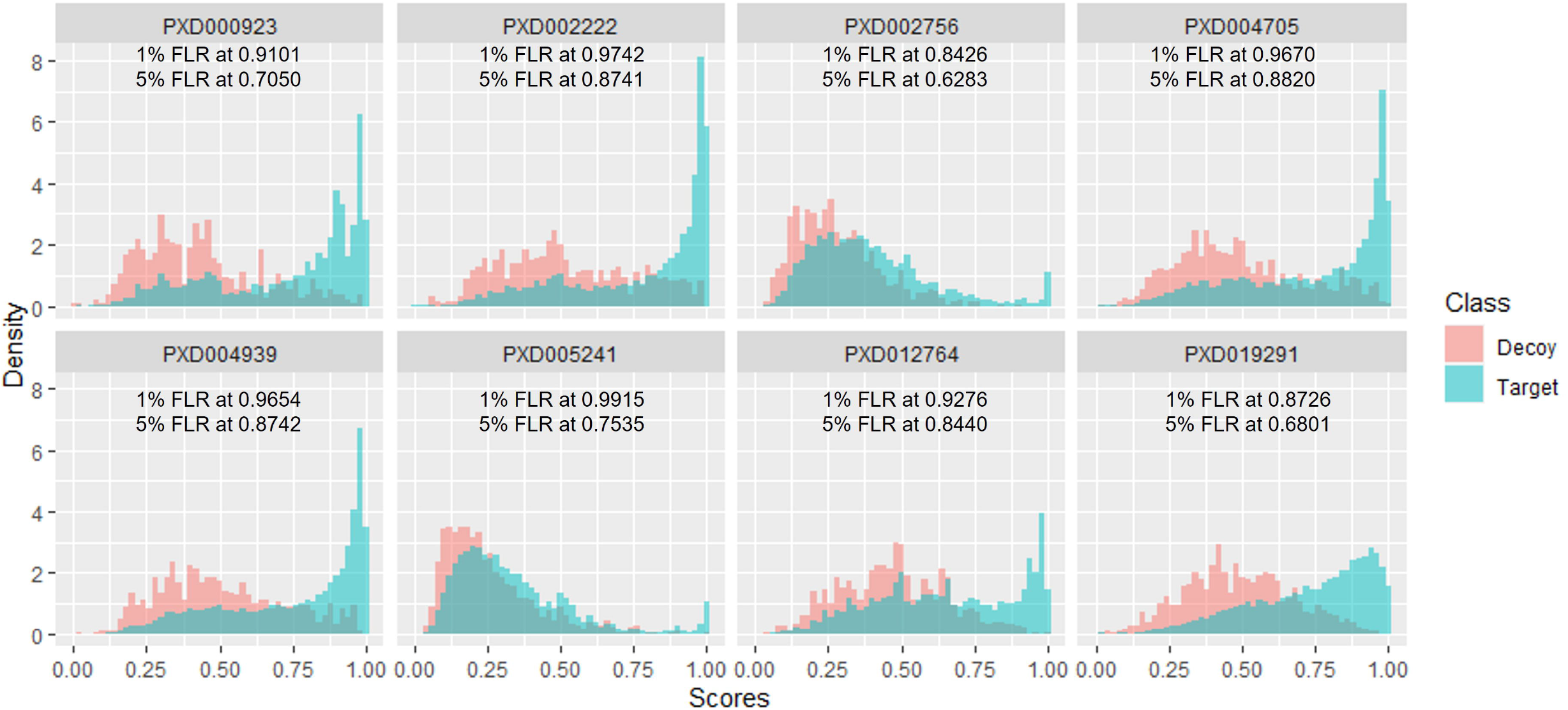
Density distributions for the number of peptide-sites assigned to decoy or target amino acids. The lowest score at 5% pAla FLR is reported below each graph.

As exemplified by the results in Table 4, the maximum method should also perform well for the data obtained using the TPP pipeline as correlated sites of a maximum score will tend to be high scoring sites as well. However, while using an MM approach seems conceptually more appropriate for at least these results generated with TPP (PTMProphet), this conclusion cannot be extended to other search engines and analysis pipelines. For the PD pipeline the MM approach seems to yield significantly better results, especially at 1% FLR threshold. Hence, for TPP users, our results suggest that the traditional approach consisting of taking the maximum score across PSM-sites linked to a peptidoform-site is acceptable and should yield similar results than those using the PSM as the basic independent unit. While PD results seems to favour by taking the mean score across a PSM before collapsing at peptidoform level.

### Collating information from independent studies

The post-processing method suggested up to this point is expected to provide scores at peptidoform-site level for a study carried out under specific experimental conditions. However, researchers may require more than one independent study to extract information of PTMs under different conditions and/or they are gathering information from publicly available data sets. How this kind of “meta-analysis” could be performed while maintaining a notion of the overall FLR could be challenging as scores from independent studies are not likely to be comparable. We illustrate the assessment of results from independent studies by collating the results from eight publicly available data sets investigating phosphorylation in rice. Table 5 displays unique protein sites for the rice data sets from data after binomial adjustment at peptidoform-site level and using traditional scoring thresholds of 0.99 and 0.95. The data shows that arbitrary thresholds are likely to underestimate or overestimate the number of high quality identifications. In this case, even the lower threshold at 0.95 will miss a significant number of overall sites (1,681 unique sites) which would be included by using the decoy approach at the most stringent criterion of 1% FLR. Although thresholding underestimates the number of confident identifications for most data sets in Table 5, it also shows that thresholding could be inflating false positive rates for some data sets like PXD002756 and PXD005241.

**Table 5.** Frequency of unique protein sites (UniqPS) after scores’ binomial adjustment at peptidoform-site level achieved by taking the maximum. Frequency of UPS for 8 rice data sets as total frequency and applying 1% and 5% pAla FLR thresholds.

| Data set | Unique sites Peptidoform-site level |  |  |  |  |
| --- | --- | --- | --- | --- | --- |
|  | All | pAla FLR =< 5% | pAla FLR =< 1% | Score<br>=>0.99 | Score<br>=>0.95 |
| PXD000923 | 4625 | 2729 | 1297 | 316 | 1379 |
| PXD002222 | 4049 | 1980 | 957 | 271 | 1005 |
| PXD002756 | 4930 | 604 | 234 | 128 | 173 |
| PXD004705 | 6507 | 2753 | 1462 | 247 | 1019 |
| PXD004939 | 6426 | 2806 | 1431 | 264 | 1072 |
| PXD005241 | 13718 | 630 | 248 | 254 | 415 |
| PXD012764 | 2607 | 820 | 616 | 69 | 336 |
| PXD019291 | 12650 | 8096 | 4302 | 549 | 1934 |
| ALL | 27951 | 11828 | 6368 | 1240 | 4687 |

At this point, when assessing information across data sets, scores are not especially informative. This is due to experimental and analytical reasons which will define a relationship between target and decoy matches specific to that study. This can be clearly observed in Figure 5. The distribution of incorrect matches (decoy) to the unknown (targets) is remarkably different across rice data sets. There are at least two completely different relationships, for PXD002756 and PXD005241 decoy and target distributions heavily overlap making it difficult to differentiate between correct and incorrect matches. The proportion of sites at 1% and 5% FLR is lower for these sets than for the other data sets. The second relationship would be preferable (as seen in PXD000923, PXD002222, PXD004705, PXD004939, PXD012764 or even PXD019291), where target high scores group on the right side of the graph forming a peak with the most confident sites while decoy matches will gather towards the left side of the graph as scores decrease. By using the decoy matches distributions across data sets as a reference, it can be observed that a same score should be interpreted as different levels of uncertainty depending on the study they come from i.e., it demonstrates that the scores are not probabilistic and/or directly comparable across studies.

A possible approach to homogenise scores could be normalisation by taking a reference across all sets such as 5% or 10% FLR. However, the method could assume that the scores are equivalent at the chosen threshold but not at the highest score as a score close to 1 could indicate different levels of uncertainty depending on the data set. Hence, although scores cannot be used for meta-analysis, FLR thresholds calculated for each data set will remain useful as they retain false positives proportionally when merging data sets with a same cut off FLR threshold.

In addition to simply taking the matches above a common threshold, further criteria based on those thresholds could be introduced to indicate different level of confidence among those identifications. For example, it is reasonable to think, that we should have increased confidence on those sites which have been identified in independent analyses.

In our example there are 11,828 unique protein sites at 5% FLR and of those 6,368 were identified at 1% FLR peptide-sequence-site level base on the Combined Algorithm (Table 5). We could further differentiate uncertainty by choosing a qualitative approach based on how many times sites are observed in independent analyses. For rice data sets we chose to label the new uncertainty groups as Gold, Silver and Bronze. Where Gold are sites observed in a minimum of 2 data sets at 1% pAla FLR; Silver as those sites observed in 1 data set at 1% pAla FLR; and Bronze any other site with pAla FLR<5%.

Results from this classification are displayed on Figure 6. Only 5 matches to alanine are returned by the Gold class suggesting a pAla FLR below 1% (0.0054), 33 matches to alanine in the Silver suggesting FLR below 2% (0.018) for that group and, 296 in the Bronze category with nearly 13% FLR (0.129) (all calculations use a STY:A ratio of 2.262 for normalisation of counts to FLR).

**Figure 6.**
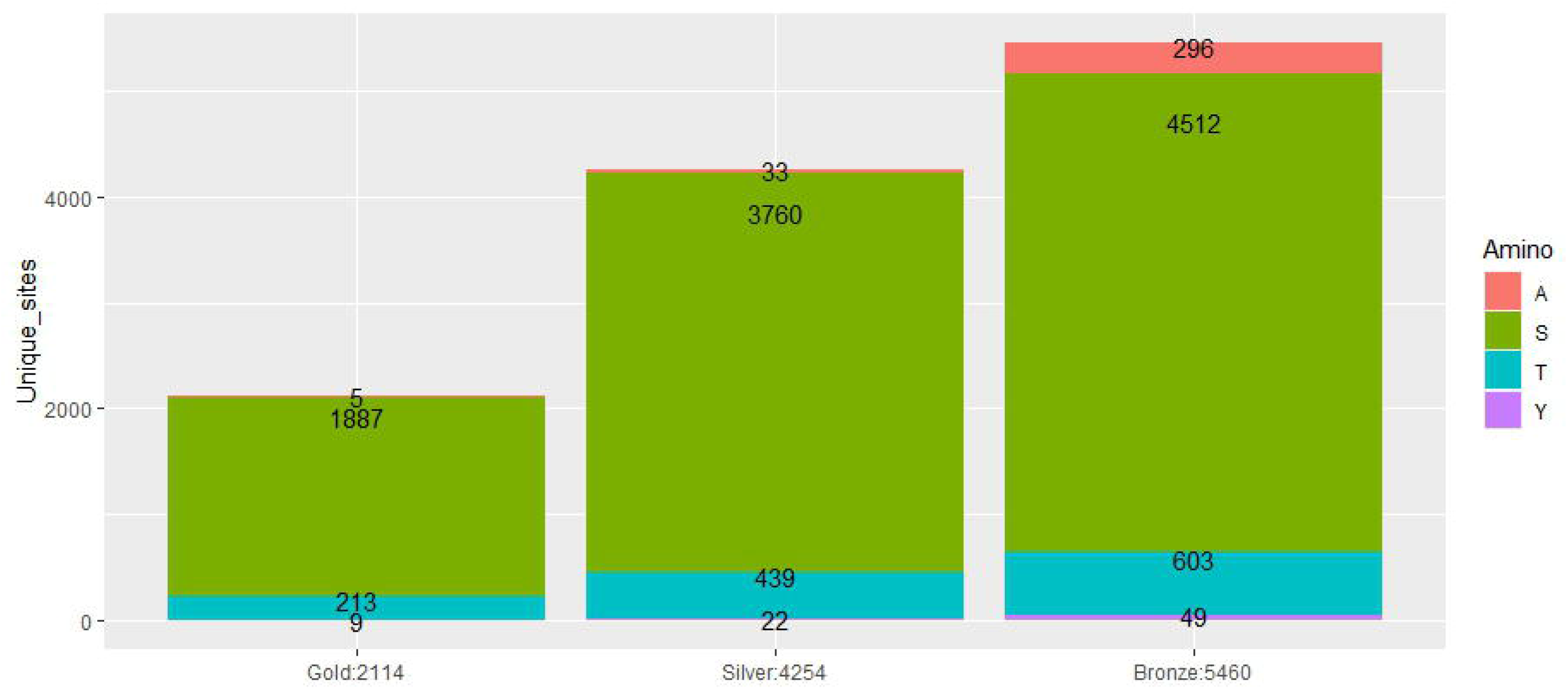
Unique protein sites obtained from 8 rice data sets with sites below 5% pAla FLR at peptide-site level by phosphorylated amino acid.

Results applying these simple criteria seem to return satisfactory level of decoy matches for our example using rice data sets. However, the relationship between decoy matches and thresholding may vary depending on the nature of the species and data being investigated and therefore criteria for gold-silver-bronze may need to be learnt on a case by case basis, e.g. optimising how many data sets are required for promotion of a site to Gold standard.

## Discussion

There is a wide range of scoring algorithms used for PTM identification and localisation. It is common practice for PTM score thresholds to be applied for real data set, which have been calibrated based on analyses of synthetic data sets for which the correct matches are known. However, it can be questioned whether these scores are truly probabilistic, hence limiting interpretability of scores across data sets. Additionally, in the context of PTM localisation, synthetic data sets may not provide a good representation of natural data sets and therefore PTM scoring calibration efforts based on synthetic data sets are not likely to be entirely reliable. These two reasons make questionable the use of a fixed scoring threshold across studies. As an alternative to scoring thresholds, it has been suggested the introduction of decoy amino acids in addition to target modifiable amino acids as variable modifications in search engines to enable calculation of data set specific FLR statistics. This approach should provide objective comparable results between independent studies based on global FLR thresholds.

Results from searches using decoy amino acids can yield many thousands of PSMs with high level of duplication and ambiguity which could hinder their interpretation. In this paper, we suggest a postprocessing workflow for LC-MS/MS based PTM localisation studies using decoy amino acids. Our workflow has simplicity at its core in which for each step simple methods are chosen to enable implementation with any basic statistical software. The workflow is data and search engine agnostic favouring exchange of findings from different sources and methodologies as long as there is a direct correlation between scores and the confidence of a PTM site being correctly identified. We suggest three steps, as follows.

First, we aim to mitigate the effect of spurious random matches, by using the binomial distribution to adjust PTM scores, with scores □ [0,1]. This adjustment takes into account differences in abundance for different peptides in a way that all scores are penalised but more heavily for sites which are observed less frequently. Our results suggested that data obtained via the TPP pipeline could benefit more from this adjustment than data extracted using the PD pipeline.

Second, we investigated simple approaches for collapsing from PSM level to Peptidoform level. Traditionally, the most common collapsing approach has been using the maximum score across all PSM-sites belonging to a peptidoform-site, even if the scores for a peptidoform with multiple PTMs come from different PSMs. This would be conceptually correct if scores for PSM-sites within a PSM are considered to be independent from each other. However, at least for the scoring methods used in this paper, PSM sites scores do not seem to be independent and the PSM average score would be a more suitable statistic than the independent scores when comparing scores across PSMs. However, given that in highly correlated scores within a PSM, to have a maximum score will lead to other PTM sites to have high scores as well. For TPP, we showed that these two approaches would lead to similar results at least for the TPP pipeline while bigger differences were observed for the PD pipeline. Therefore, the traditional approach of taking the maximum score would be preferred in general as it would perform well for PSM level with correlated and independent scores while those using the pipelines with PD at its core may obtain better performance by using the mean PSM score.

Third, we have also shown that scores from independent studies were not comparable as they are likely to be closely linked to the experimental and data collection procedures of each study. Therefore, we suggested a simple approach to combining evidence from independent studies. By retaining the matches to decoy PTM sites based of different FLR thresholds overall FLR estimates of combined data can be determined.

A limitation to the results presented here is that these are based on two analysis pipelines and scoring methods. Further analyses would be required to demonstrate that the simple postprocessing approaches explained above would also perform well for data acquisition via other analysis pipelines and PTM scoring algorithms. This is especially important for those scores ∉ [0,1] for which an additional step might be required.

We believe our FLR estimates to be broad FLR indicators due to the complex nature of PTM scoring. We have shown that synthetic data sets often used to calibrate scoring methods are not likely to be representative of natural data sets and therefore it is not possible to fully ascertain performance but enables comparisons with current practice.

## Supporting information

Analysis of PD data sets

## Funding

We gratefully acknowledge funding from BBSRC that contributed to this work [BB/S017054/1, BB/T019557/1].

## Data availability

All original mass-spectrometry data can be found http://proteomecentral.proteomexchange.org/ as:

- PXD000138: http://proteomecentral.proteomexchange.org/cgi/GetDataset?ID=PXD000138
- PXD000612: http://proteomecentral.proteomexchange.org/cgi/GetDataset?ID=PXD000612
- PXD000923: http://proteomecentral.proteomexchange.org/cgi/GetDataset?ID=PXD000923
- PXD002222: http://proteomecentral.proteomexchange.org/cgi/GetDataset?ID=PXD002222
- PXD002756: http://proteomecentral.proteomexchange.org/cgi/GetDataset?ID=PXD002756
- PXD004705: http://proteomecentral.proteomexchange.org/cgi/GetDataset?ID=PXD004705
- PXD004939: http://proteomecentral.proteomexchange.org/cgi/GetDataset?ID=PXD004939
- PXD005241: http://proteomecentral.proteomexchange.org/cgi/GetDataset?ID=PXD005241
- PXD007058: http://proteomecentral.proteomexchange.org/cgi/GetDataset?ID=PXD007058
- PXD008355: http://proteomecentral.proteomexchange.org/cgi/GetDataset?ID=PXD008355
- PXD012764: http://proteomecentral.proteomexchange.org/cgi/GetDataset?ID=PXD012764
- PXD019291: http://proteomecentral.proteomexchange.org/cgi/GetDataset?ID=PXD019291

Analysed data at PSM-site level can be found as supplementary data.

R code used to produce all tables and figures in this report can be found at <u>omcamacho/pSTYA_post.processing (github.com)</u>

## Notes

https://github.com/omcamacho/pSTYA_post.processing

http://proteomecentral.proteomexchange.org/

### Competing Interest Statement

The authors have declared no competing interest.

