## Supplementary material for "Assessing multiple evidence streams to decide on confidence for identification of post-translational modifications, within and across data sets": Analysis of PD data sets

SUPPLEMENTARY INFORMATION

Supp Table 1. Parameters used for peptide identification and PTM site localisation using TPP.

| Data set ProteomeXchange Unique ID | **Peptide Mass Tolerance**  **(ppm)** | **Fragment Bin Tolerance (Da)** | **Digest Mode** | **Max Missed Cleavages** | **Fixed Mods** | **Variable Mods** | **Max Variable PTMs** |
| --- | --- | --- | --- | --- | --- | --- | --- |
| PXD000138 | 5 | 0.02 | Tryptic | 4 | Carbamidomethylation (C) | Oxidation (M),  Phospho (STYA),  N-terminal acetylation,  Ammonia loss (QC),  Pyro-Glu (EQ on the N-terminus),  Deamination (NQ) | 5 |
| PXD007058 | 20 | 0.02 | Tryptic | 4 | Carbamidomethylation (C) | Oxidation (MWP),  Phospho (STYA),  Pyrophospho (STY)¶,  N-terminal acetylation,  Ammonia loss (QC),  Pyro-Glu (EQ on the N-terminus),  Deamination (NQ) | 5 |
| PXD008355 | 7 | 0.02 | Tryptic | 2 | Carbamidomethylation (C) | Oxidation (M),  Phospho (STYA),  N-terminal acetylation,  Ammonia loss (QC),  Pyro-Glu (EQ on the N-terminus),  Deamination (NQ) | 5 |
| PXD000612 | 7 | 0.02 | Tryptic | 2 | Carbamidomethylation (C) | Oxidation (M),  Phospho (STYA),  N-terminal acetylation,  Ammonia loss (QC),  Pyro-Glu (EQ on the N-terminus),  Deamination (NQ) | 5 |
| PXD000923 | 50 | 0.02 | Tryptic | 2 | Carbamidomethylation (C) | Oxidation(MW)  N-terminal acetylation, ammonia loss (QC), pyro-glu (E), deamination (NQ) and phosphorylation (STYA) | 5 |
| PXD002222 | 20 | 0.02 | Tryptic | 2 | Carbamidomethylation (C) | Oxidation(MW)  N-terminal acetylation ammonia loss (QC), pyro-glu (E), deamination (NQ) and phosphorylation (STYA) | 5 |
| PXD002756 | 20 | 1.0005 | Tryptic | 2 | Carbamidomethylation (C) | Oxidation(MW)  N-terminal acetylation ammonia loss (QC), pyro-glu (E), deamination (NQ) and phosphorylation (STYA) | 5 |
| PXD004705 | 20 | 0.02 | Tryptic | 2 | Carbamidomethylation (C) | Oxidation(MW)  N-terminal acetylation ammonia loss (QC), pyro-glu (E), deamination (NQ) and phosphorylation (STYA) | 5 |
| PXD004939 | 20 | 0.02 | Tryptic | 2 | Carbamidomethylation (C) | Oxidation(MW)  N-terminal acetylation ammonia loss (QC), pyro-glu (E), deamination (NQ) and phosphorylation (STYA) | 5 |
| PXD005241 | 20 | 1.0005 | Tryptic | 2 | Carbamidomethylation (C) | Oxidation(MW)  N-terminal acetylation ammonia loss (QC), pyro-glu (E), deamination (NQ) and phosphorylation (STYA) | 5 |
| PXD012764 | 10 | 0.02 | Tryptic | 2 | Carbamidomethylation (C)  iTRAX8plex label | Oxidation(MW)  N-terminal acetylation ammonia loss (QC), pyro-glu (E), deamination (NQ) and phosphorylation (STYA) | 5 |
| PXD019291 | 10 | 0.02 | Tryptic | 2 | Carbamidomethylation (C) | Oxidation(MW)  N-terminal acetylation ammonia loss (QC), pyro-glu (E), deamination (NQ) and phosphorylation (STYA) | 5 |

Supp Table 2. Parameters used for peptide identification and PTM site localisation using Mascot/ptmRS.

|  | Peptide Mass Tolerance | Fragment Bin Tolerance | Digest Mode | Max missed cleavages | Fixed Mods | Variable Mods | Max Variable PTMs |
| --- | --- | --- | --- | --- | --- | --- | --- |
| PXD007058 (Synthetic data set) | 20.0 ppm | 0.02 Da | Tryptic | 2 | Carbamidomethylation (C) | Oxidation (MWP)  Phospho (STYA)  Pyrophospho (STY) | 3 |
| PXD008355 (Arabidopsis data set) | 10.0 ppm | 0.02 Da | Tryptic | 2 | Carbamidomethylation (C) | Oxidation (M)  Phospho (STYA)  N-terminal acetylation  Ammonia loss (QC)  Pyro-Glu (EQ on the Nterminus)  Deamination (NQ) | 3 |

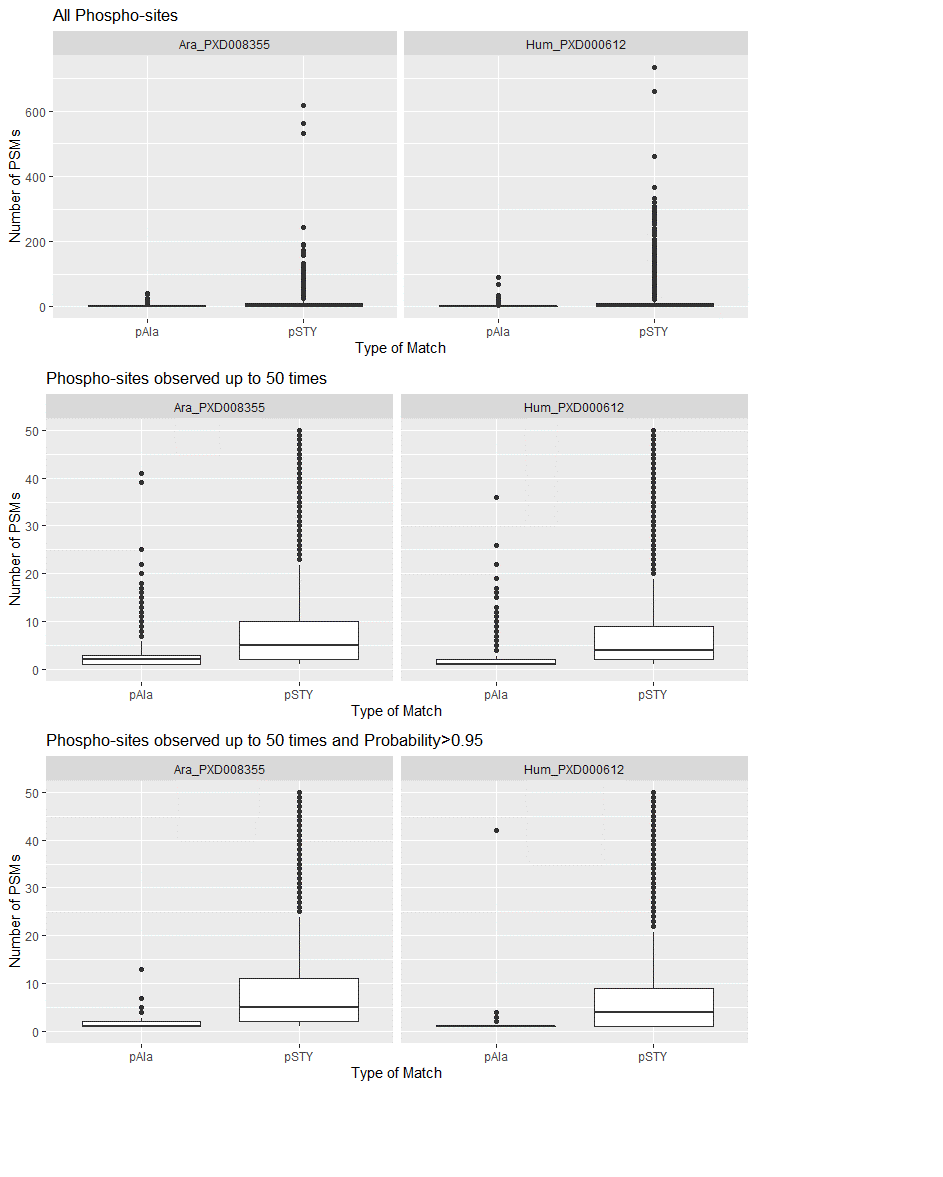

Supp Figure 1. Number of PSMs matching specific sites by type of match for Human and Arabidopsis natural data sets analysed using the PD pipeline. pAla matches correspond to decoy phospho-sites and pSTY target matches. Top 10 panels display all data, middle 10 panels display phospho-sites observed up to 50 times and, the bottom 10 panels are those phospho-sites with scores probability above 0.95 and observed up to 50 times.

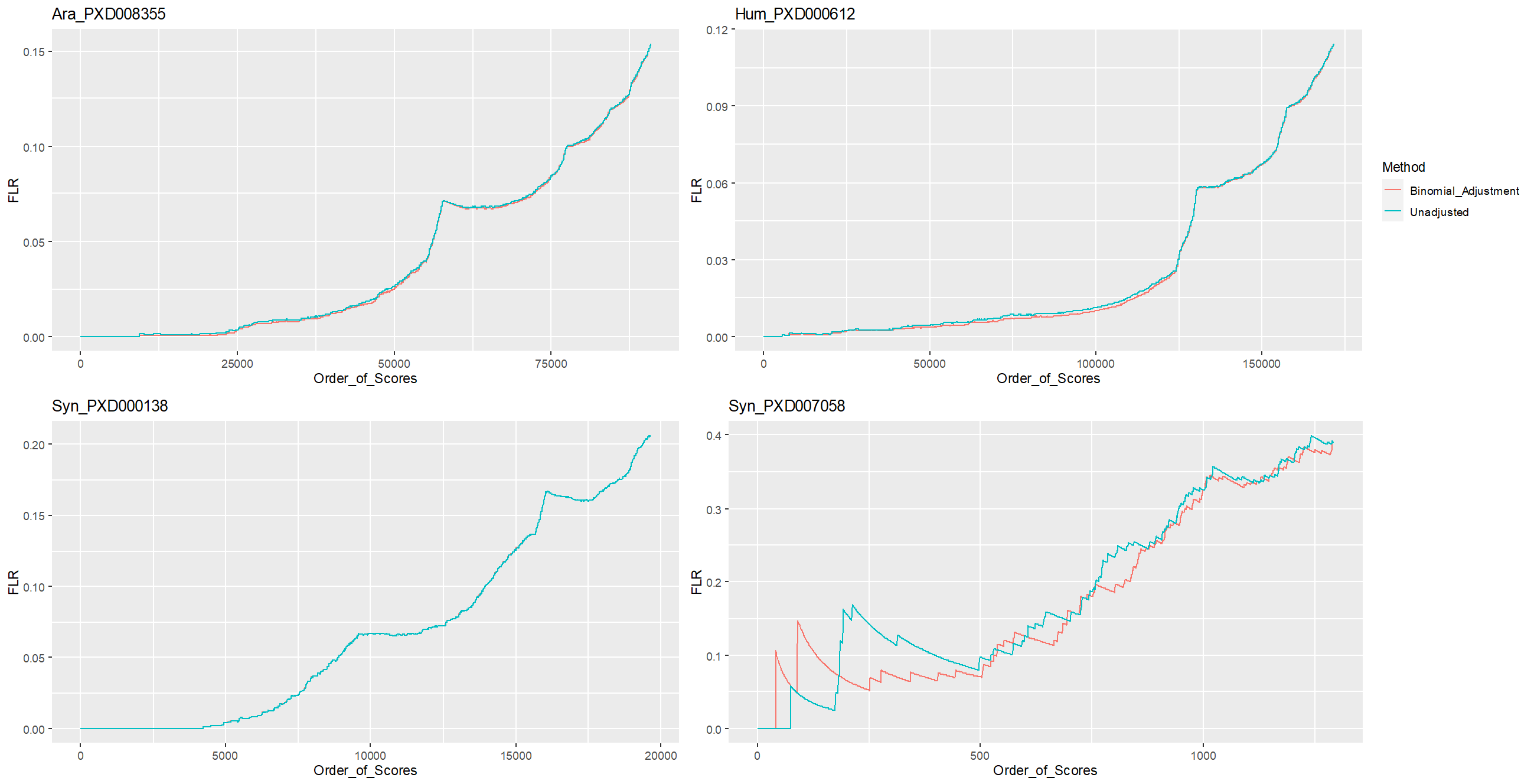

Supp Figure 2. pAla FLR at PSM-site level by ordered unadjusted scores (Blue), using the Product_Adjustment (Green) and, using the Bi-nomial_Adjustment (Red). Data sets analysed using PD pipeline.

Supp Table 3. Number of PSM-site matches for Arabidopsis, Human and two synthetic data sets analysed using the PD pipeline and thresholds at 1%, 2.5%, 5% and 10% pAla FLR based on unadjusted results and adjusted using the Product_Adjusted and Binomial Adjusted approaches. For the synthetic data sets, the Real_FLR is also displayed, calculated as the proportion of pAla PSM-site matches at each pAla FLR threshold.

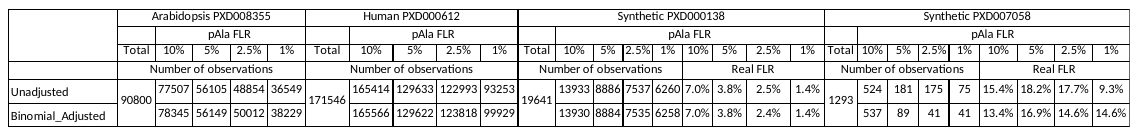

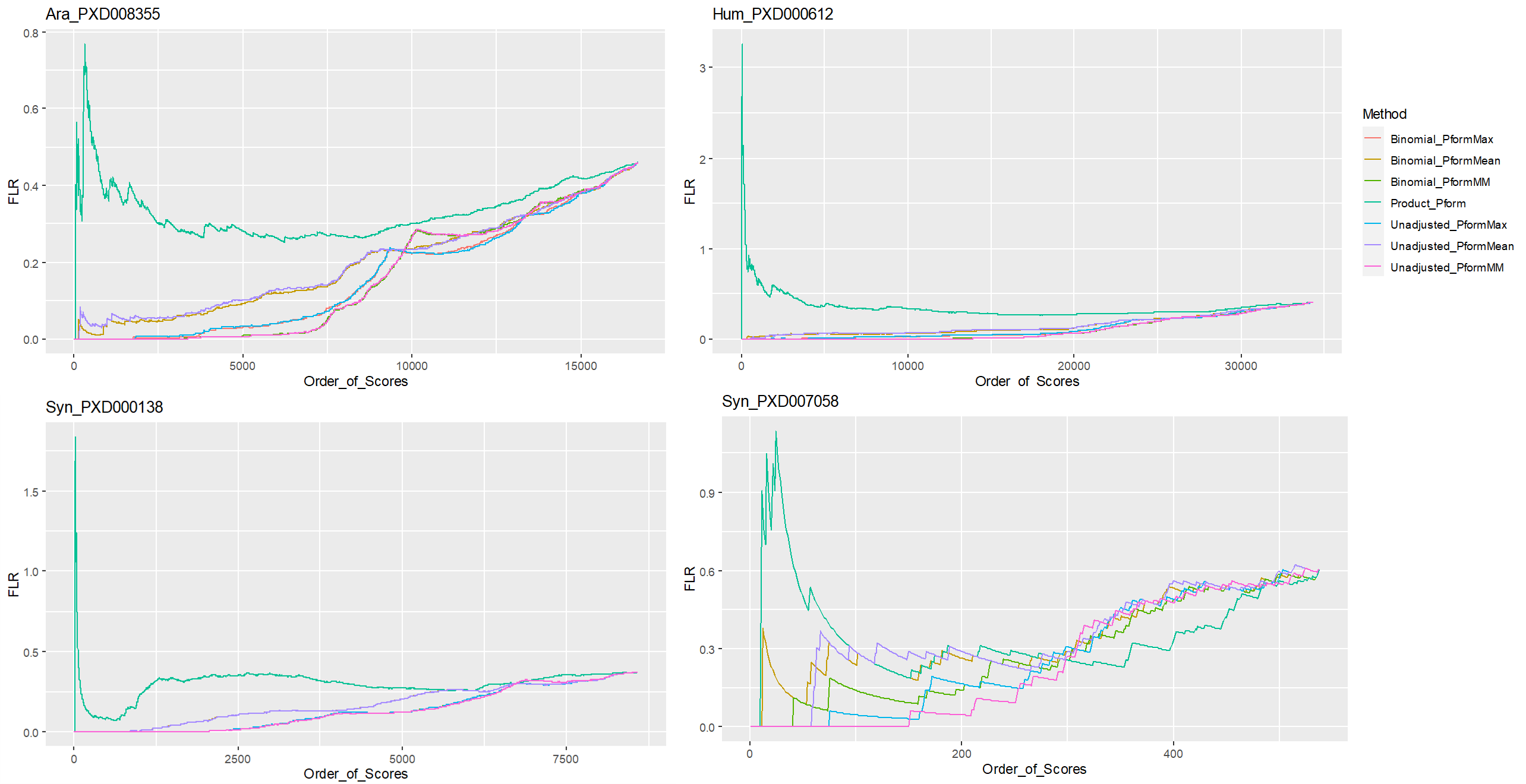

Supp Figure 3. pAla FLR at Peptidoform-site level by ordered scores for six collapsing approaches in two natural data sets and two synthetic. Data analysed using the PD pipeline.

Supp Table 4. Data sets analysed using the PD pipeline. Number of phospho-sites at Peptidoform-site level for six collapsing approaches. Results are reported overall and for pAla 1%, 5% and 10% FLR thresholds. Real FLR is also displayed for the synthetic data sets.

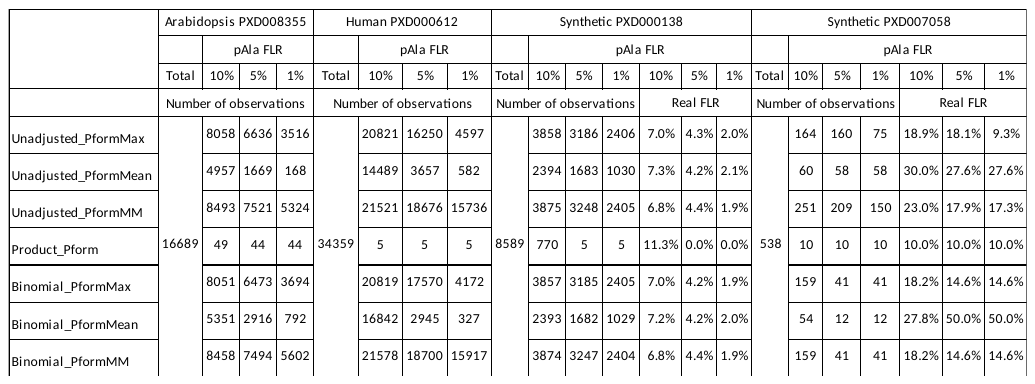
